# One-shot design elevates functional expression levels of a voltage-gated potassium channel

**DOI:** 10.1101/2022.12.28.522065

**Authors:** Jonathan Jacob Weinstein, Chandamita Saikia, Izhar Karbat, Adi Goldenzweig, Eitan Reuveny, Sarel Jacob Fleishman

## Abstract

Membrane proteins play critical physiological roles as receptors, channels, pumps, and transporters. Despite their importance, however, low expression levels often hamper the experimental characterization of membrane proteins. We present an automated and web-accessible design algorithm called mPROSS (https://mPROSS.weizmann.ac.il), which uses phylogenetic analysis and an atomistic potential, including an empirical lipophilicity scale, to improve native-state energy. As a stringent test, we apply mPROSS to the Kv1.2-Kv2.1 paddle chimera voltage-gated potassium channel. Four designs, encoding 9-26 mutations relative to the parental channel, were functional and maintained potassium-selective permeation and voltage dependence in *Xenopus* oocytes with up to 14-fold increase in whole-cell current densities. Additionally, single-channel recordings reveal no significant change in the channel-opening probability nor in unitary conductance, indicating that functional expression levels increase without impacting the activity profile of individual channels. Our results suggest that the expression levels of other dynamical channels and receptors may be enhanced through one-shot design calculations.

**Significance statement:** Heterologous expression levels of membrane proteins are often low, limiting research and applications. We combine homologous-sequence analysis with Rosetta atomistic calculations to enable one-shot design of dozens of mutations that improve native-state energy. Applied to a voltage-gated potassium channel, designs exhibited up to 14-fold improved functional expression levels in oocytes with almost no change in the single-channel activity profile. This design approach may accelerate research of many challenging membrane proteins, including receptors, channels, and transporters.

## Introduction

Membrane proteins (MPs) are gatekeepers to the cell with critical roles in signal transduction and metabolism. Despite their importance, MPs often exhibit marginal stability and low functional expression levels (1). The functional expression of MPs in the eukaryotic plasma membrane depends on their stability (2) as well as on a multilayered trafficking and quality-control system embedded in the endoplasmic reticulum and the Golgi apparatus (1). MPs may be mislocalized or targeted for degradation at any stage of biogenesis due to signals that are only partly understood (1).

Methods for improving MP functional expression have had a transformative impact on research but required intense experimental effort. For instance, breakthroughs in determining the structure and function of voltage-gated potassium (Kv) channels and G-protein coupled receptors demanded iterative mutagenesis and experimental screening to increase heterologous expression levels (3–7). Other approaches for improving MP expressibility included deep mutational scanning and genetic recombination of natural homologs(8–10), but these approaches demand medium to high-throughput screening which is impractical for most MPs(7). Computational design methods have also been developed, but their reliability has so far been limited, demanding multiple iterations of design and expeirment(11–15).

Here, we ask whether the heterologous expression levels of a MP can be improved through one-shot design calculations that improve native-state energy. Recently, our lab developed the PROSS algorithm for increasing the stability and heterologous expression levels of water-soluble proteins(16, 17). PROSS combines phylogenetic analysis and Rosetta atomistic calculations to design dozens of mutations that stabilize the native state. In a benchmark, PROSS improved thermal stability or expressibility in most proteins and required testing fewer than five constructs per target(18). In some cases, PROSS improved thermal stability by ≥20 °C and boosted heterologous expression levels by several orders of magnitude without compromising activity(16). PROSS has been applied successfully to optimize therapeutic enzymes(19), vaccine immunogens(20, 21), soluble receptors(22), and recently, dramatically increased the expression levels of previously uncharacterized enzymes starting from AI-based model structures(23). At its core, however, PROSS uses a water-soluble energy function that is inappropriate for MP design. To develop a membrane version of PROSS (mPROSS), we changed the energy function to one that accounts for the membrane environment(24) by using the dsTβL empirical scale of amino acid insertion into the plasma membrane(25, 26). Furthermore, the new energy function addresses both water-soluble and membrane domains to account for the large extra-membrane domains often observed in MPs.

As a stringent test, we apply mPROSS to a model Kv channel, the paddle chimera of Kv1.2-Kv2.1(27). We chose this synthetic construct as it is one of the only Kv channels characterized by crystallographic analysis at high resolution(27). Furthermore, the paddle chimera exhibits 20-fold lower K^+^ current densities than the wild-type Kv1.2, suggesting that its functional expression levels may be impaired(28). Kv channels are highly selective, mostly homotetrameric channels found throughout the animal kingdom(29). They have important roles in the physiology of many organs and in various autoimmune and cardiac muscular and neurological pathologies(30, 31). Moreover, Kv channels are highly dynamic, undergoing large conformational rearrangements that control channel opening in response to changes in the electric potential across the plasma membrane. Many natural and synthetic toxins target ion channels, and the paddle chimera is an important model for investigating these interactions(32). Furthermore, Kv channels can be investigated at the single-molecule level, providing detailed information on functionality that complements bulk experiments. We tested four mPROSS designs encoding 9 to 26 mutations relative to the parental channel and found that they exhibited higher expressibility in *Xenopus* oocytes and increased conductance while preserving voltage sensitivity and channel-opening probability and conductance. The results demonstrate that automated design calculations can be applied even to a large, oligomeric, dynamic, and structurally complex MP to improve its heterologous functional expression levels without impacting its functional properties. Moreover, the results suggest that the premisses that underlie successful stability design in soluble proteins(33) may extend to MPs. mPROSS can be applied, in principle, to any MP through a webserver that is freely available to academic users (https://mPROSS.weizmann.ac.il).

## Results

### Improving native-state energy in membrane proteins

mPROSS combines phylogenetic analysis with Rosetta atomistic calculations to design variants that exhibit improved native-state energy. Starting from the amino acid sequence of the target MP, we search the non-redundant (nr) sequence database(34) for homologs and at each position, eliminate mutations that are rarely observed in the natural diversity. The protein is then embedded in a virtual membrane(35, 36) using the Orientations of Proteins in Membranes (OPM) algorithm(37) and relaxed in Rosetta. Next, we use Rosetta atomistic design calculations to eliminate destabilizing single-point mutations (Supplementary Table 1). Last, we use Rosetta combinatorial sequence design to compute several low-energy multipoint mutants that may incorporate dozens of mutations relative to the parental protein (Fig. 1a). In all atomistic design calculations, regions that are critical for activity are immutable (Supplementary Table 3). Furthermore, we disallow mutations at positions that exhibit Gly, Pro, or positions immediately preceding Pro, because they may have roles in determining MP structure and dynamics(38).

**Figure 1.**
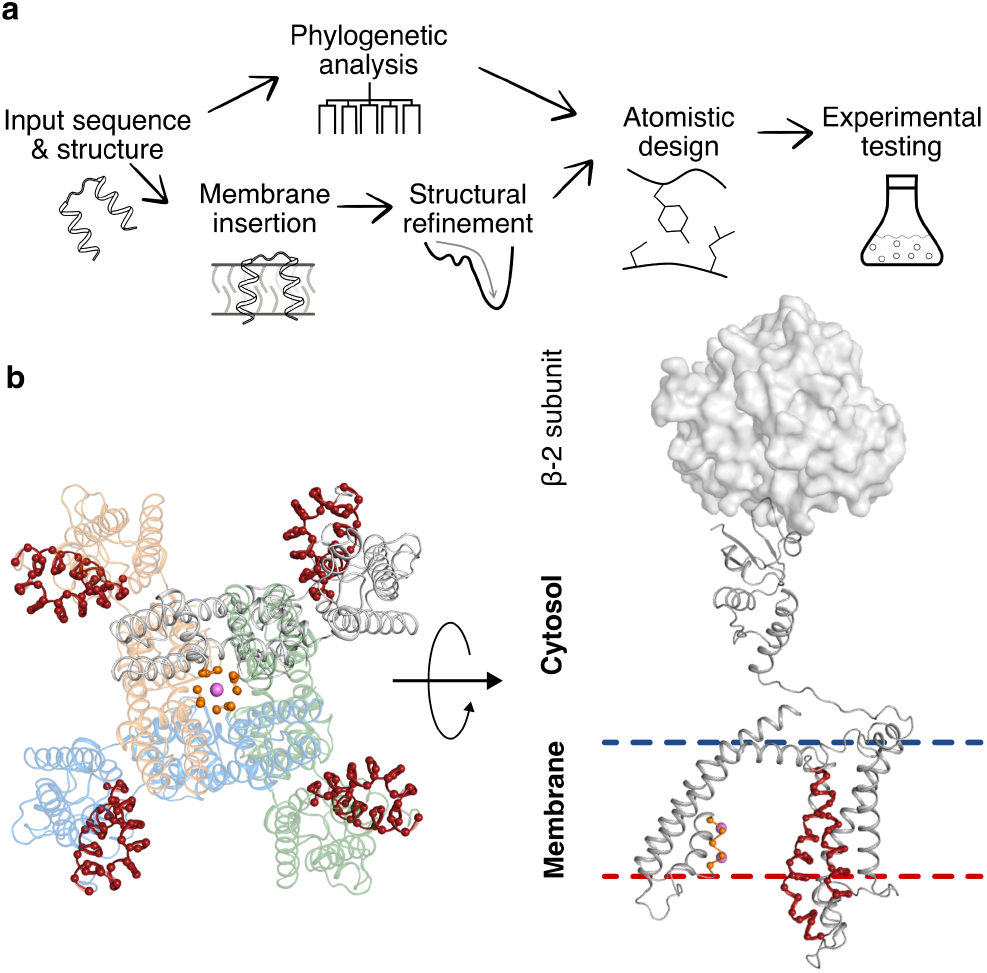
Steps in applying mPROSS to the Kv channel. (a) Scheme of the mPROSS algorithm. mPROSS starts with a phylogenetic analysis of homologs of the target protein and inserts the protein into a virtual membrane. The structure is minimized and subjected to combinatorial design calculations, generating designs with different mutational loads for experimental analysis. (b) Overview of the paddle chimera structure, including the β-2 soluble subunits responsible for attenuating channel dynamics (PDB entry: 2R9R). The selectivity filter is indicated in orange, and the voltage sensor is in red. The left-hand side shows the tetrameric arrangement of the channel looking from the extracellular domain toward the fourfold symmetry axis of the channel. Pink spheres indicate K^+^ ions. The right-hand side shows a side view of one protomer, and a single β-2 subunit is shown as a molecular surface.

To account for the physical heterogeneity of the membrane environment, mPROSS uses the ref2015_memb energy function in all atomistic calculations(24). In the water-soluble regions, ref2015_memb is identical to the standard ref2015 energy function(24, 39), whereas within the membrane, it recapitulates the experimentally determined MP insertion scale dsTβL(24, 25). The dsTβL amino acid insertion scale exhibits membrane-depth-dependent lipophilicity. For instance, the hydrophobic amino acids Leu, Ile, and Phe are highly favored near the membrane midplane. In contrast, polar amino acids, such as Asn and Asp, are disfavored in the membrane, as are the helix-distorting amino acids, Gly and Pro. The basic amino acids Arg and Lys exhibit asymmetric potentials, favoring their localization to the membrane inner leaflet, following the “positive-inside” rule(40). We recently demonstrated that ref2015_memb accurately predicts structures of homomeric single-pass MPs *ab initio(24)* and is effective in *de novo* design of receptor-like transmembrane domains of defined oligomeric state and geometry(41). Here, we extend it to design low-energy variants of large and complex MPs.

As a benchmark, we applied mPROSS to 20 nonredundant (<80% sequence identity(42)) multi-spanning membrane proteins (Fig. 2, Supplementary Table 2). These proteins range in size from 103 to 492 amino acids and span multiple functional classes, including channels, transporters, enzymes, and receptors.

**Figure 2.**
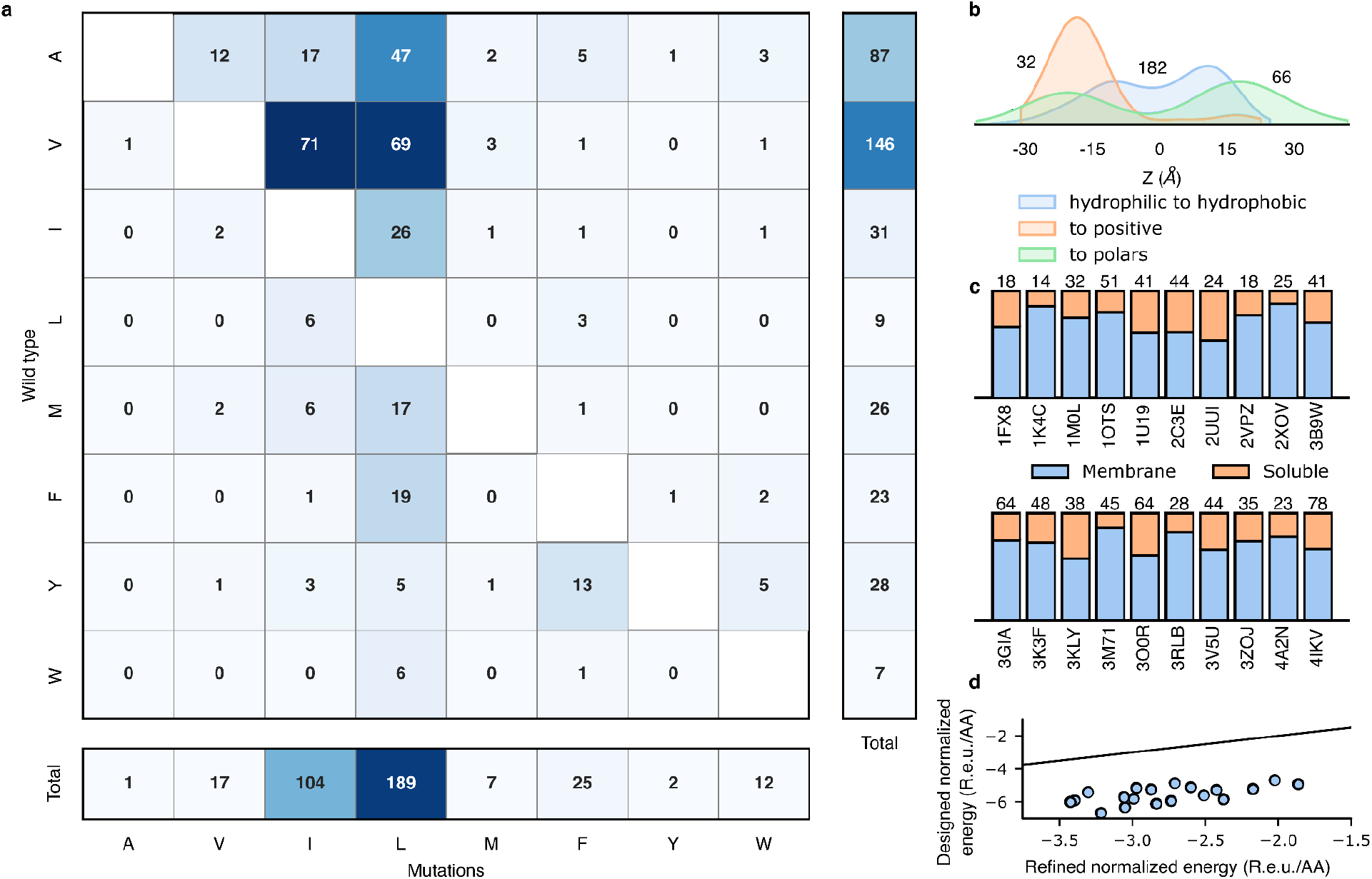
mPROSS introduces mutations that increase lipophilicity and the “positive-inside” rule. For each of the 20 proteins in the benchmark, we analyzed a single mPROSS design that incorporates approximately 15% mutations, the maximal fraction of mutations we recommend for testing. (**A**) The number of hydrophobic-to-hydrophobic mutations within the membrane domain. (**B**) Normalized distributions of mutations across the membrane. The horizontal axis represents the distance to the membrane midplane, with negative and positive values representing the intra- and extracellular domains, respectively. Numbers denote the number of mutations in each distribution. (**C**) Fraction of mutations in the membrane or soluble domain in each protein. The number of mutations is noted above the bars, and Protein Data Bank (PDB) entries are noted below the bars. (**D**) Energies of refined and designed proteins. Line marks *x*=*y*. R.e.u./AA denotes the total Rosetta energy divided by protein length.

Near the membrane midplane, mPROSS introduces predominantly hydrophobic-to-hydrophobic and hydrophilic-to-hydrophobic mutations, and mutations to aromatic identities (51, 30, and 9%, respectively, Fig. 2a). Moreover, most mutations increase side chain lipophilicity, mainly by introducing Leu, the most lipophilic amino acid(25). In contrast, mutations in the soluble domain are of a mixed character, as expected (Fig. 2b & c, Supplementary Fig. 1). Furthermore, only a small fraction of mutations in the membrane are to polar residues (7%), mostly at positions buried within the protein where they would be shielded from lipid and predominantly to the mildly polar Thr. Strikingly, mutations to the positive Arg and Lys identities mostly occur in the inner-membrane leaflet or intracellular domain (90%), following the “positive-inside” rule (Fig. 2b)(40, 43, 44). The overall effect of introducing dozens of mPROSS mutations is a significant improvement in the Rosetta energy relative to the parental protein (Fig. 2d). We conclude that mutations introduced by mPROSS may improve membrane localization and enforce its native topology.

### Paddle chimera designs exhibit high conductance levels

To avoid mutations that impact channel function, we disabled design in the voltage-sensor domain (S3 and S4, Fig. 1b), the K^+^-selectivity filter, and the inner vestibule. Additionally, we restricted positions in the tetrameric interfaces to maintain the oligomeric structure (Supplementary Table 3). mPROSS automatically generates 18 designs, of which we chose four that exhibit a large number of mutations (designs 12, 16, 17, and 18; these were renumbered for clarity in the remainder of the manuscript to D1-D4, respectively). Following visual inspection, we removed up to ten mutations from each design (Supplementary Tables 3 & 4). The four designs introduced 9-26 mutations (5-13% of mutable positions; Fig. 3 and Supplementary File 2) and were subjected to experimental analysis. We note that as in other applications of PROSS (16), the Rosetta atomistic energies correlate with the number of designed mutations (Table S4) but may not directly correlate with stability or expression levels. Because Rosetta energies do not provide an accurate prediction of expressibility, we typically test experimentally 3-5 design, as we did here.

**Figure 3.**
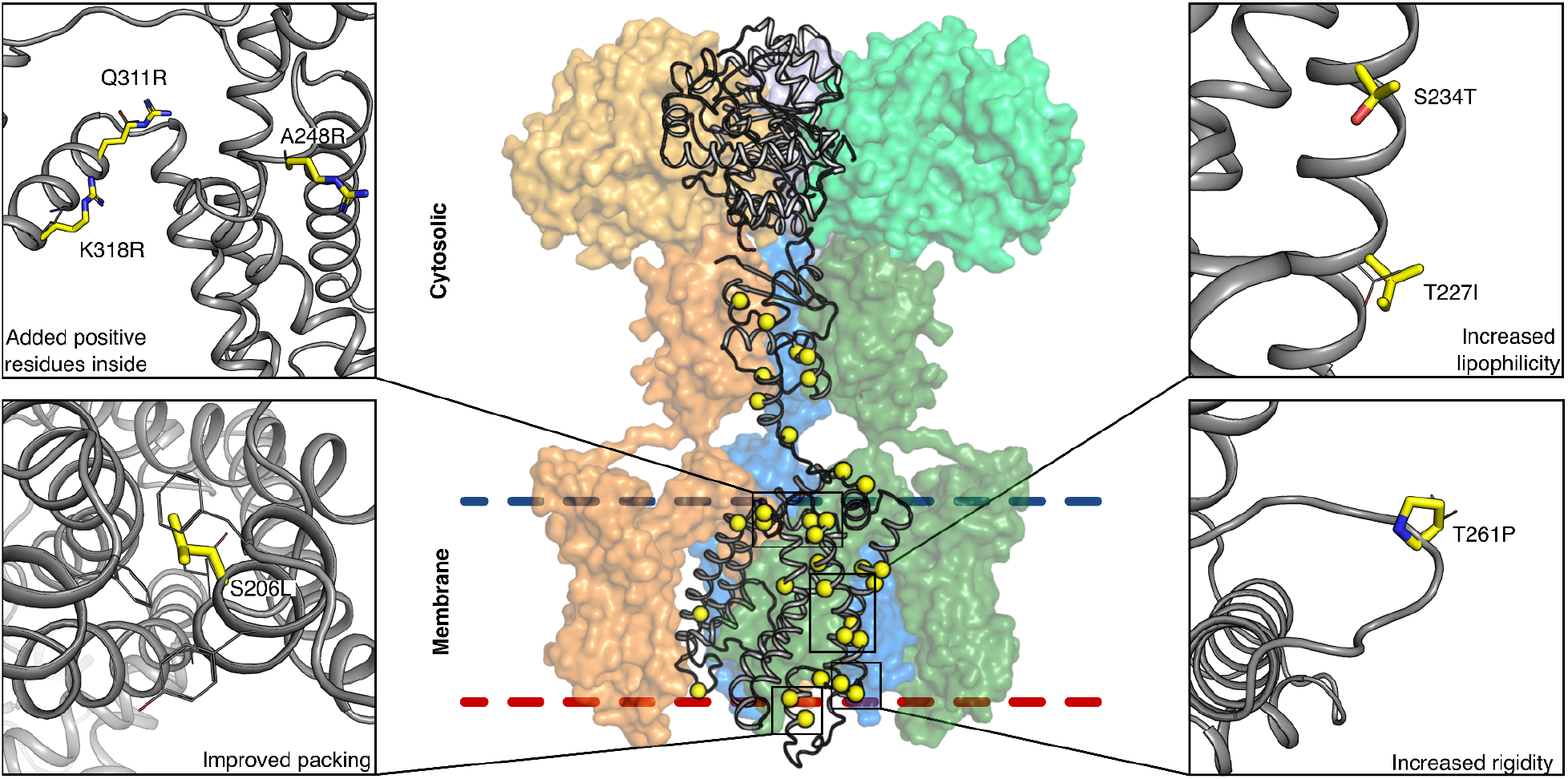
Designed mutations improve protein lipophilicity and native-state molecular interactions. mPROSS designs both the membrane and cytosolic domains. Mutations increase the lipophilicity of membrane-exposed positions, add positively charged residues in the inner leaflet (following the “positive-inside” rule(40)), rigidify loops, and improve interhelix packing. The protein mainchain is shown in gray, and mutations are shown in thick yellow sticks or spheres. Only one protomer is shown in cartoon representation, and the other three are represented as colored molecular surfaces. The β-2 subunits are shown in lighter colors (top).

To determine the yield of functional channels, we recorded channel activity and characterized voltage dependence of the parental channel and four designs in *Xenopus laevis* oocytes using a single voltage step from a holding potential of -80 to -20 mV for 120 ms. Remarkably, all designs exhibited higher current amplitudes than the parental channel (Fig. 4a), and designs D1 and D2 exhibited significantly elevated amplitudes (5- and 14-fold relative to the paddle chimera, respectively). Because designs D3 and D4 conducted currents similar to D2 (Supplementary Table 6), we focused our subsequent analysis on D1 and D2.

**Figure 4:**
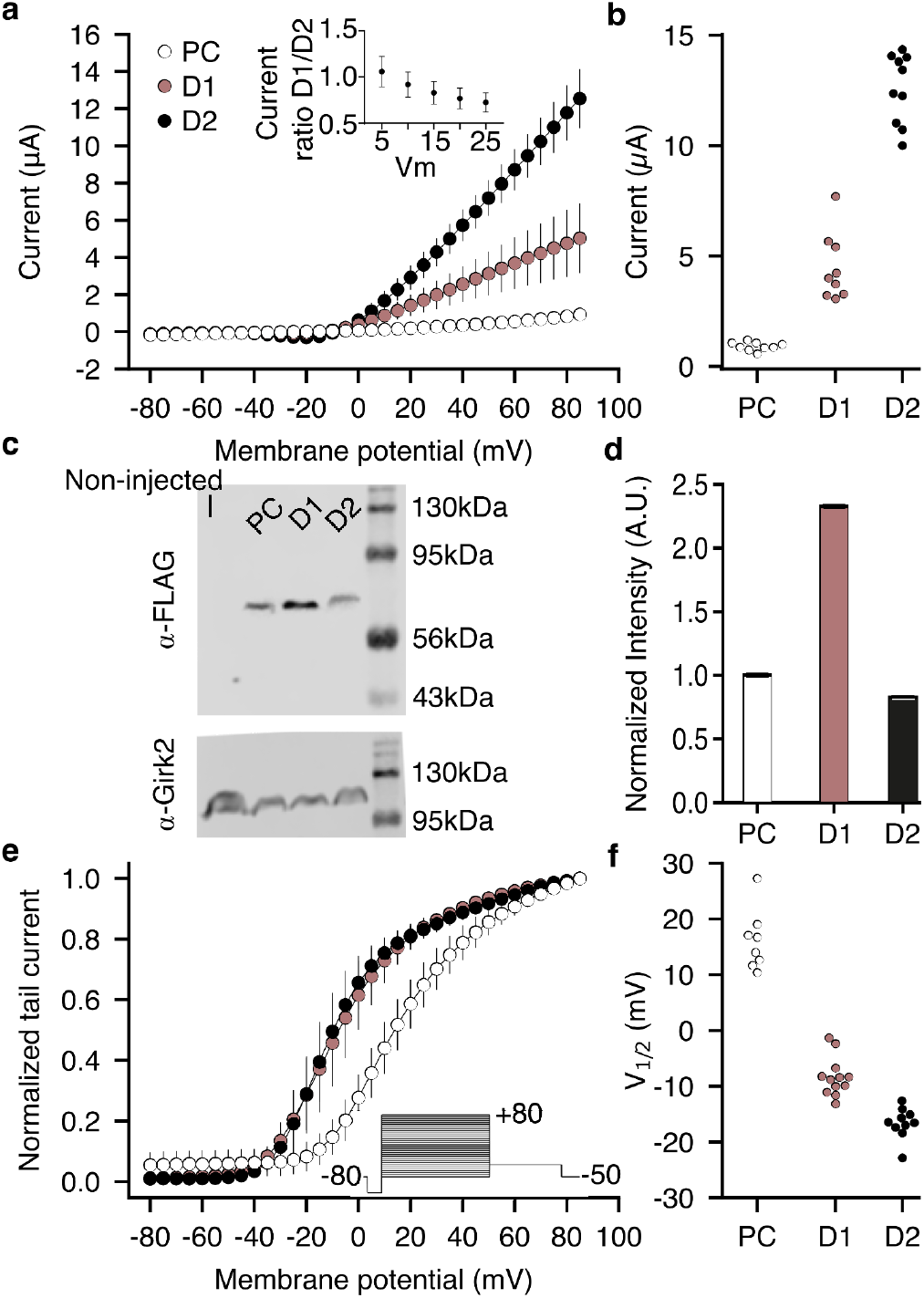
Current-voltage relationship of the Kv1.2-Kv2.1 paddle chimera and its designs. (a) Currents of the paddle chimera (PC) and designs are plotted against varying test voltages. Each curve represents recordings from *n*⩾8 oocytes. Error bars represent standard deviations. (inset) Ratios of currents observed for D1 and D2 at voltages in the range 5-25 mV. (b) Average currents (paddle chimera: 0.91±0.19 µA, D1: 4.65±1.52 µA, D2: 12.50±1.63 µA). (c) Western blot analysis of total membrane expression. The blots were labeled with anti-FLAG (top) or with anti-GIRK2 (as control; bottom) antibodies. (d) Normalized blot intensities. The experiment was performed in triplicate. Error bars represent the standard deviation. (e) Normalized tail currents were recorded at -50 mV as a function of the test pulse voltage (see inset). Measured tail current amplitudes were fitted with a single-component Boltzmann equation (solid lines), from which V_1/2_ of activation was inferred. (f) Inferred V_1/2_ values (paddle chimera: 16.10±5.39 mV, D1: -8.33±3.49 mV, and, D2: -16.61±2.75 mV).

We sought to understand the large difference in current amplitudes between D1 and D2 using mutational analysis. The two designs differ by six mutations, precluding a complete analysis of all combinations of mutations and prompting us to nominate mutations based on structural analysis. We noticed that the mutation Thr271Arg is present in both designs and faces the membrane inner leaflet (all position numbers according to Long et al.(27) unless indicated otherwise). We hypothesized that the designed Arg could interact with the phospholipid head groups to stabilize the open state. To test this hypothesis, we reverted the mutation Thr271Arg on the D2 background. The revertant exhibited voltage sensitivity similar to D2, however, refuting a substantial role for this mutation in stabilizing the open state (Supplementary Fig. 2 and Supplementary Table 5). We then examined several other mutations that differ between D1 and D2 that we deemed likely to affect conductance. On the D2 background, we individually reverted Phe261Leu and Val280Leu (within the voltage-sensing domain), Met340Leu (on the S4-S5 linker), and Tyr151Phe, Arg133Lys and Glu75Asp (within the cytosolic tetramerization domain). Remarkably, none of the revertants recapitulated D1 currents, and some had currents higher than D2. These single substitution analyses indicate that the increased current in D2 likely results from the interaction of multiple designed mutations. They also highlight a strength of the mPROSS strategy of designing multipoint mutations by demonstrating that, individually, mutations may exhibit only a limited impact on expressibility(33).

The increased currents we observed for all designs can stem from higher plasma membrane expression levels, elevated single-channel conductance, or higher opening probability. First, we estimated total expression levels from immunoblots of oocyte total membrane fractions, finding that D1 expressed at approximately twofold higher levels than the parental channel (Fig. 4c & d), in agreement with its increased whole-cell currents. By contrast, D2 exhibited similar expression levels to the parental channel (Fig. 4c & d), suggesting that increased total expression levels are not the reason for the higher current amplitudes observed for this design. Furthermore, at potentials where the opening probability is lower than maximum, the currents observed for D1 and D2 were not significantly different (Fig 4a, inset). This suggests that the difference in current density between D1 and D2 may be due to a change in the number of functional channels at the plasma membrane. We then tested whether the increased current amplitudes arise from a hyperpolarizing shift in the voltage dependence of the channels. We recorded currents elicited by voltage steps from -80mV to +80mV in 90mM KCl-containing solution and determined the relative conductance from the resulting tail currents at -50mV. D1 and D2 exhibited only a slight shift in voltage dependence relative to the parental protein (Fig. 4e & f). Nevertheless, the current amplitudes of the designed channels were higher than those of the paddle chimera across the entire voltage range, including at positive potentials where the opening probability is maximal. These data suggest that the shift in voltage dependence is not the main cause for the apparent increase in current amplitudes observed for the designs.

Finally, we tested whether the increased currents were due to a change in the unitary conductance or the channel opening probability. We recorded single-channel currents under the cell-attached configuration of the patch-clamp technique in oocytes. Single-channel conductance for the parental channel and D2 were similar (10.4±0.4 and 9.7±2.3 pS, respectively; Fig. 5). Furthermore, no change was observed in the opening probability between these channels, with Po of wild type and D2 of 0.009+/-0.004 and 0.011+/-0.005 at 0mV, respectively. Taken together, these results rule out the possibility that the designed mutations increase potassium fluxes or stabilize the channel open state and suggest, again, that the increase in ion conductance is due to a higher abundance of functional channels in the plasma membrane.

**Figure 5.**
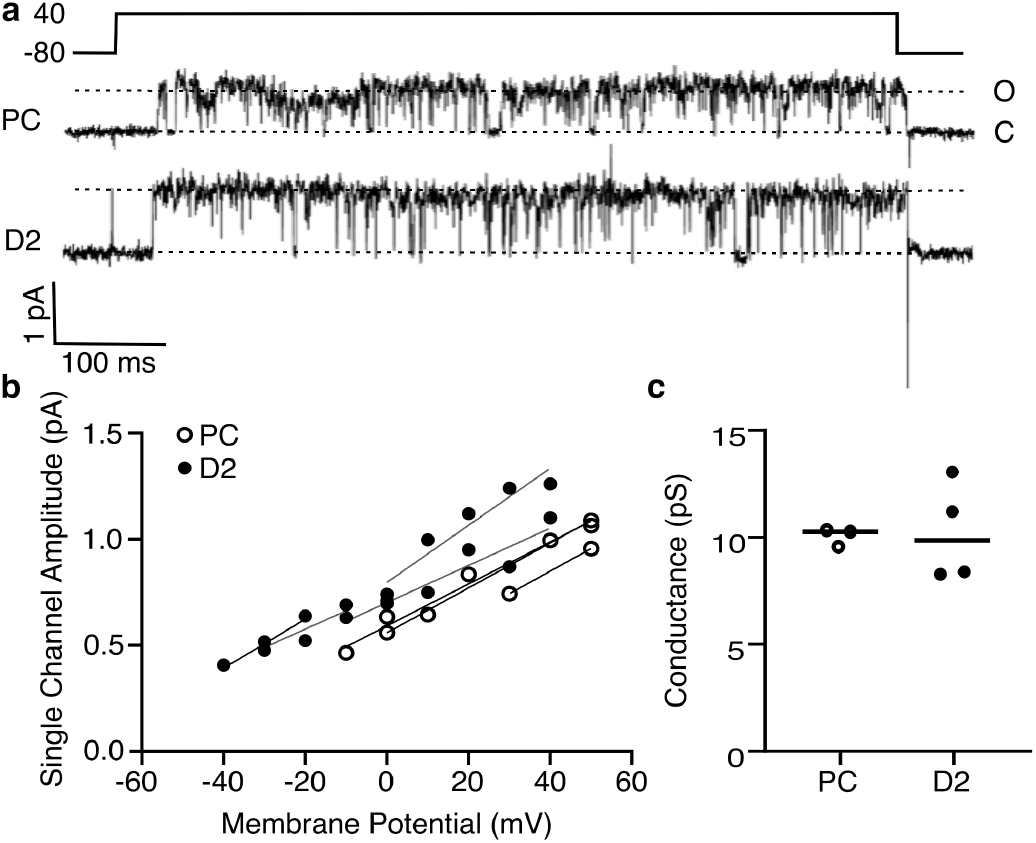
Design D2 behaves similarly to the parental channel at the single-channel level. (a) Representative current traces were elicited at +40 mV from a holding potential of -80 mV under cell-attached configuration in *Xenopus* oocytes for the parental channel chimera and D2. The upward deflection indicates the open state. (b) Individual single-channel amplitudes at different potentials from all recorded patches. (c) Channel conductance derived from the current to voltage slopes shown in (b).

## Discussion

Eukaryotic MP biogenesis is a complex process that involves multiple quality-control steps(1). Despite this complexity, our results provide a case study that validates a one-shot computational design approach that introduces dozens of mutations selected only according to native-state energy and phylogenetic propensity. Some designs exhibit substantially higher functional expression levels than the parental protein without exhibiting observable activity differences at the single-protein level. Increased functional expression may be due to a higher level of plasma membrane localization or to a greater fraction of natively folded and active channels by stabilizing interactions with membrane lipids or within the designs. It is also possible that the designs alter the protein folding pathway, limiting off-pathway misfolding and slowing the kinetics of unfolding(33). Further experimental analysis will be required to decide between these possibilities. In a companion paper, we demonstrate that mPROSS can also increase stability and expression levels while retaining wild-type levels of activity in a human membrane enzyme based on its AlphaFold2 model structure(45). The combination of AlphaFold2 and mPROSS extends MP design to proteins that have not been successfully subjected to experimental structure determination. Taken together with the broad success of the PROSS algorithm in designing stable and expressible soluble proteins (16, 18–20, 46), we conclude that protein expression may depend, to a large extent, on native-state energy, whether in soluble or membrane-embedded proteins.

## Acknowledgements

We thank Nir Fluman for critical discussions. Research in the Fleishman lab was supported by the Israel Science Foundation (1844), the European Research Council through a Consolidator Award (815379), the Dr. Barry Sherman Institute for Medicinal Chemistry, and a donation in memory of Sam Switzer. Research in the Reuveny lab was supported by the Israeli Science Foundation (349/22) and the Wilner Family Fund. ER is the incumbent of the Charles H. Hollenberg Professorial Chair.

## Methods

### mPROSS algorithm

mPROSS is identical to PROSS2, except where explicitly mentioned.

#### Phylogenetic analysis

The phylogenetic analysis is identical to PROSS2(17). Briefly, sequences are collected using BLASTP(47, 48) against the non-redundant database. Only hits with sequence identity >35% and coverage >70% are kept. Sequences are then aligned using MUSCLE(49) and clustered using CD-HIT(50). The MSA is then segmented by secondary structure elements. The sub-MSA for each sequence is pruned of sequences with different gaps than the query sequence. A sub-PSSM (Position Specific Scoring Matrix) is then created for each segment, and all resulting sub-PSSMs are concatenated to create a PSSM for the complete protein sequence(51). Any mutation with PSSM score <0 or absent from the MSA is eliminated from consideration. Additionally, the PSSM score is used to bias the energy function to favor mutations that are more common in the natural diversity.

#### Constraints

To avoid introducing deleterious mutations, certain positions are restricted from design. The first and last three positions of each chain, including next to missing density gaps, are automatically restricted, and all non-canonical residues are automatically restricted. If the user specifies interacting proteins or small molecule or ions ligands, all positions within 8 or 5Å are restricted, respectively. The user can also specify additional positions to restrict from design.

#### Rosetta calculations

All RosettaScripts and flags are in Supplementary File 1.

Protein structures are inserted into a virtual membrane(36) using the OPM algorithm(37). Structures are then refined using the Rosetta full-atom membrane protein energy function, ref2015_memb, and a “softened” version of ref2015_memb, where van der Waals repulsion is weakened(24) to enable crossing energy barriers. Five refined structures, and the best scoring one is used in all subsequent structural calculations.

A virtual mutational scan, where each non-restricted position is mutated to each possible amino acid, is conducted using the Rosetta FilterScan mover. Amino acids absent from the MSA in each position are avoided in the mutational scan. The virtual mutation scan applies a set of thresholds and weights (Supplementary Table 1), generating a set of mutations that pass the ΔΔ*G* threshold. Each set is considered a sequence space for combinatorial design.

The amino acid sequence of the refined structure is then optimized within each sequence space. Due to the high convergence we observe in PROSS design calculations(16), we only carry out two independent trajectories and use the best-scoring model for each sequence space.

### Experimental testing

#### Gene cloning

Gene fragments encoding the paddle chimera and designed channels, codon-optimized for expression in *Xenopus*, were ordered from Twist Biosciences. A backbone vector containing the 3’ and 5’ segments of the K_v_1.2 gene (including the UTR regions) in pUC57-Kan was ordered from Genscript. The final constructs were assembled using golden-gate cloning(52), taking advantage of *BsaI* sites engineered into the gene fragments and the backbone vector. The final constructs encode for the full-length paddle chimera with a 5’ 10-histidine tag and a thrombin cleavage site, cloned into the SacII/EcoRI sites of pUC57-Kan.

Two-Electrode Voltage Clamp cRNAs encoding for the Kv1.2-Kv2.1 paddle chimera and the various designs were transcribed using T7 mMESSAGE mMACHINE Transcription Kit (Ambion) and stored as stock solutions at −80°C. *Xenopus laevis* female frogs surgery and oocyte isolation and injection were according to a published protocol(28). Potassium currents from the injected Xenopus oocytes were measured using a two-electrode voltage clamp technique with a Gene Clamp 500 amplifier (Molecular Devices). Some measurements that required high external K^+^ were conducted in a 90K solution containing 90mM KCl, 2mM MgCl_2_, and 10mM HEPES at pH 7.4. Data were sampled at 10 kHz and filtered at 5 kHz using a Digidata 1550A device controlled by pCLAMP 10.5 (Molecular Devices). Capacitance transients and leak currents were removed by subtracting a scaled control trace utilizing a P/4 protocol.

#### Western blot

Injected oocytes were checked for expression using the two-electrode voltage-clamp technique (above). Oocytes expressing paddle chimera or designed channels were collected and lyzed using Tris buffer (pH 8) containing protease inhibitors (APExBIO, catalog No. K1007). The lyzed oocytes were manually homogenized and centrifuged at 1500 g for 15 minutes. The supernatant was collected and centrifuged again at 80,000 g for 1 hour to isolate the membrane fraction. The pellet was resuspended in lysis buffer, and protein concentration was determined using the Bradford assay. A total of 20 mg protein was separated by 12% SDS-PAGE and then transferred onto nitrocellulose paper. Membranes were incubated with mouse monoclonal anti-flag antibody (abcam, catalog No. ab125243) for 16 hours. Post-incubation, membranes were washed with Dulbecco′s phosphate-buffered saline (Merck) with an HRP-conjugated anti-mouse secondary antibody at 1:1000 (Jackson ImmunoResearch Laboratories). The blot was developed using Odyssey® XF Imaging System (LI-COR Biosciences). For the controls, anti-GIRK2 (Alomone lab, catalog No. APC-006) and HRP conjugated anti-rabbit secondary antibodies were added to label intrinsic membrane proteins expressed in Xenopus oocytes. Controls confirm that the same amount of protein was loaded in each well.

#### Single-channel recordings

Single-channel currents from devitalized injected oocytes were measured by the cell-attached patch-clamp technique using an Axopatch 200B amplifier (Molecular Devices)(53). The bath solution was 90K (see above), and the pipette solution contained ND96 supplemented with 50 μM GdCl_3_. Current traces were elicited by step depolarization pulses for the various voltages (−30 to +40) from a holding potential of -80 mV for 600 ms (Clampex 10.4, Molecular Devices), and were low-pass filtered at 1 kHz, digitized at 10 kHz (Digidata 1550A, Molecular Devices). Leak and transient currents were subtracted using current traces with no openings. Po was calculated as percent time spent in the open state at 100 steps to 0 mV. Patches that contained more that one channel were corrected accordingly. Data analysis was performed using Clampfit 11.2 (Molecular Devices) and Prism 9.4 (GraphPad).

#### Animal Handling

All procedures employed on experimental animals, including their transportation, routine care, and use in experiments, was conducted in accordance with the Israel animal welfare law and guidelines, NIH guidelines, the Animal Welfare Act, the ethical standards and guidelines of FP7, H2020 with the EU directive 86/609/EEC, as well as the revised directive 2010/63/EU on the protection of animals used for scientific purposes. Approved Weizmann Institute IACUC.

## Data availability

All data are included in the manuscript and/or supporting information.

## Supplementary information

**Supplementary Figure 1.**
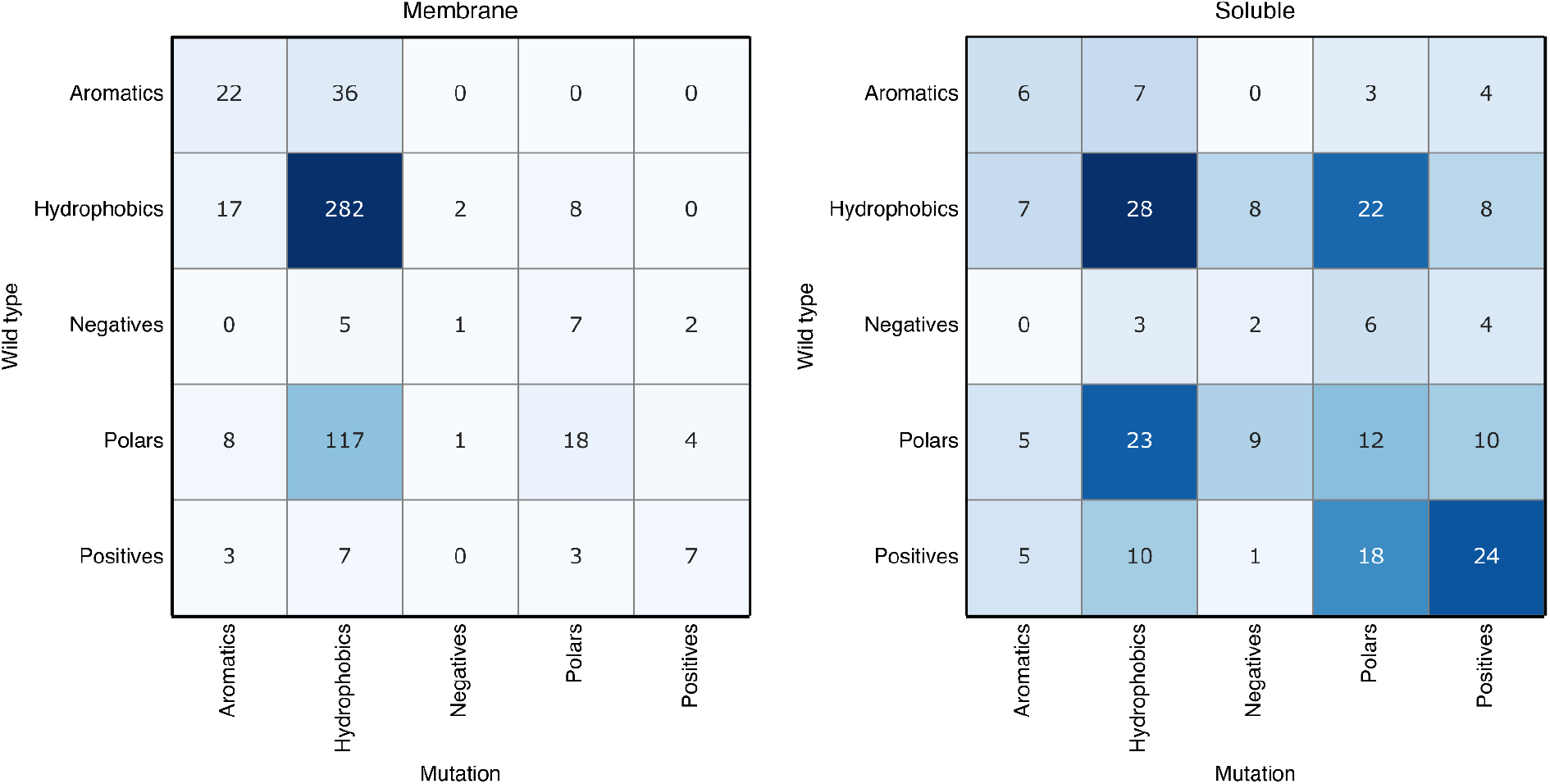
Comparing mutation types between the membrane and soluble domains. Similar to Figure 2a, the analysis is based on a single design for each protein in the benchmark that mutates approximately 15% of the protein relative to the wild type.

**Supplementary Figure 2:**
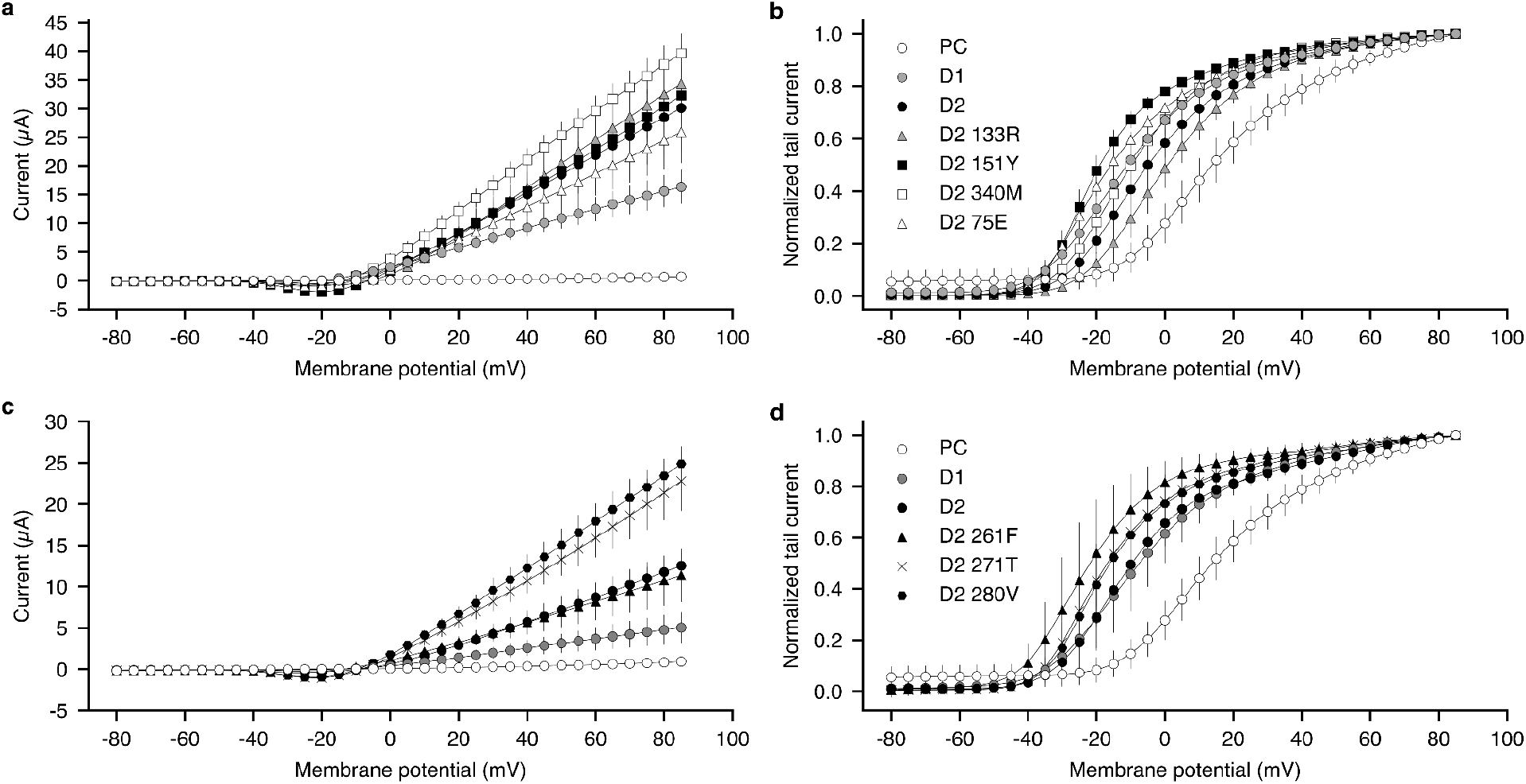
Current-voltage relationship of Kv1.2-Kv2.1 paddle chimera and designs compared to single point mutation revertants. (a & c) Currents of paddle chimera and Kv1.2-Kv2.1 paddle chimera designs are plotted against varying test voltages. Each curve represents recordings from 10 oocytes, and error bars represent the standard deviation. (b & d) Normalized tail currents were recorded at-50mV as a function of the test pulse voltage (see inset Fig. 4e). Data for paddle chimera, D1, and D2 are the same as from Fig 4. Expression was not quantified in these experiments, but equal amounts of cRNAs were injected to decrease variability.

**Supplementary Table 1.** Weights and thresholds for the computational mutational scan. ΔΔG threshold - the threshold over which mutations are discarded. PSSM threshold - PSSM score under which mutations are discarded. PSSM weight - the weight used to bias ref2015_memb energy function with the PSSM score.

| Index | $\Delta\Delta G$ threshold | PSSM threshold | PSSM Weight |
| --- | --- | --- | --- |
| 1 | -4 | 5 | 0.2 |
| 2 | -4 | 5 | 0.2 |
| 3 | -3.5 | 4.5 | 0.2 |
| 4 | -3 | 4 | 0.2 |
| 5 | -3 | 3 | 0.3 |
| 6 | -2.5 | 3 | 0.3 |
| 7 | -2.5 | 2 | 0.3 |
| 8 | -2 | 2 | 0.4 |
| 9 | -2 | 2 | 0.4 |
| 10 | -2 | 2 | 0.4 |
| 11 | -2 | 2 | 0.4 |
| 12 | -2 | 1 | 0.4 |
| 13 | -2 | 1 | 0.4 |
| 14 | -1 | 0 | 0.5 |
| 15 | -1 | 0 | 0.5 |
| 16 | -0.5 | 0 | 0.5 |
| 17 | 0 | 0 | 0.5 |
| 18 | 0.5 | 0 | 0.5 |

**Supplementary Table 2.**
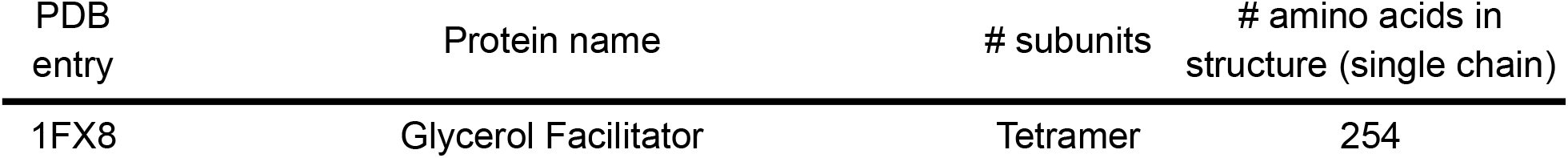
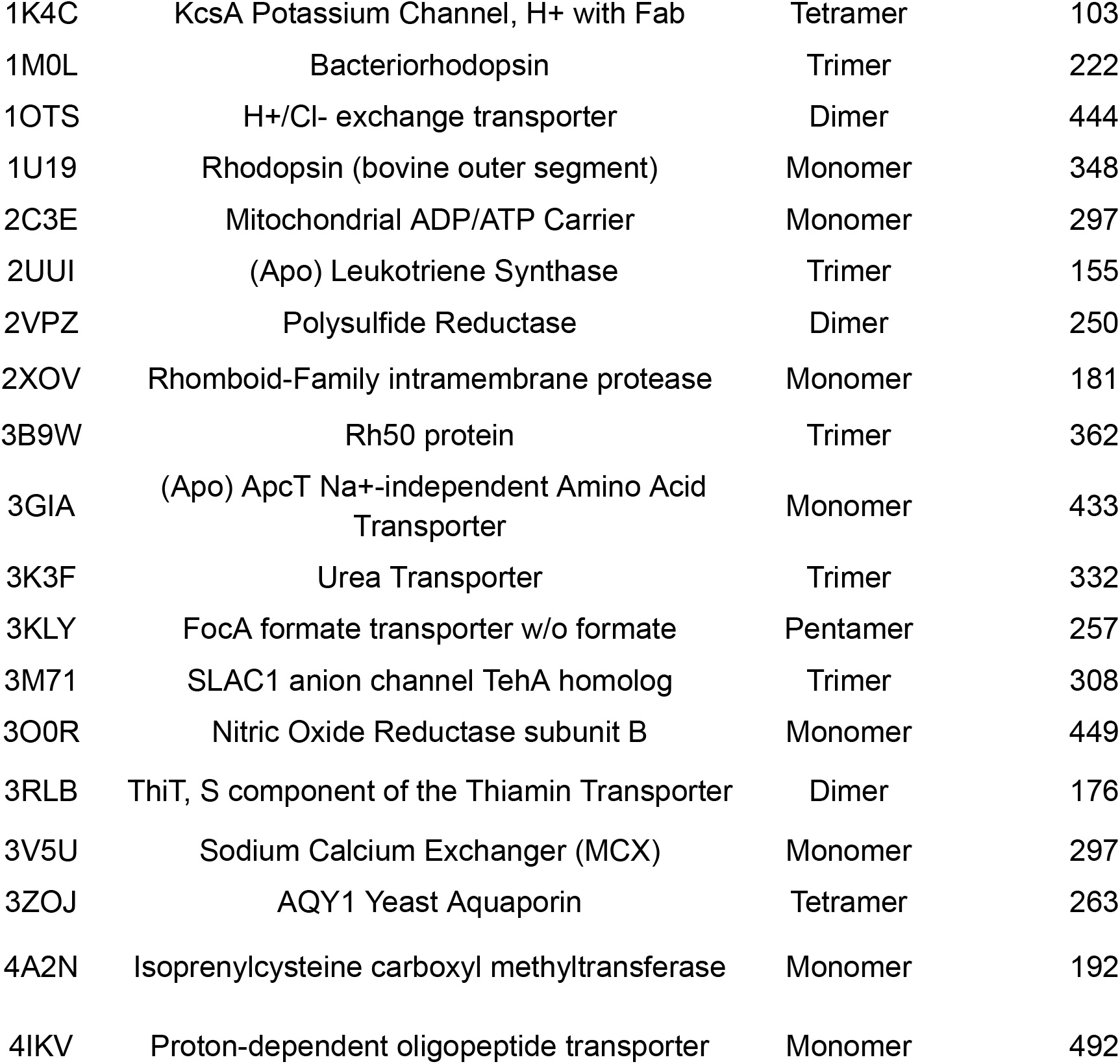
Protein structures used in the benchmark.

**Supplementary Table 3.**
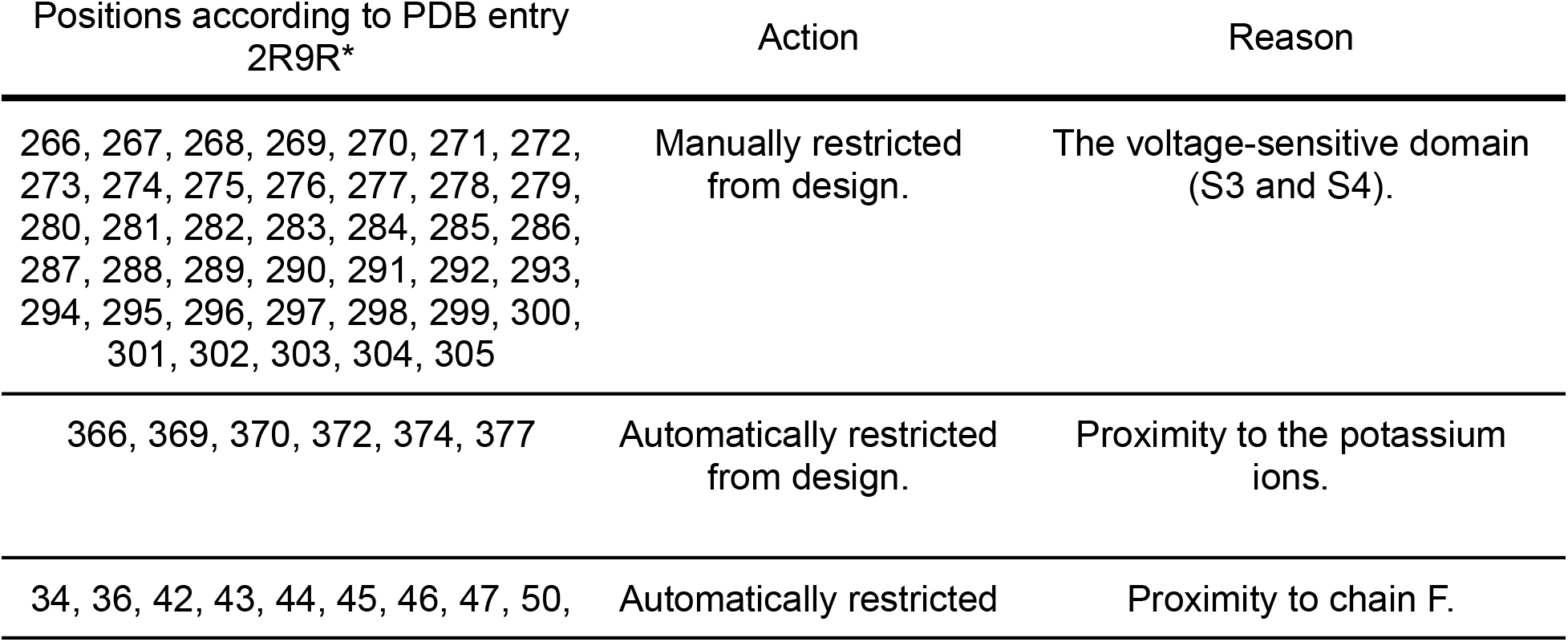
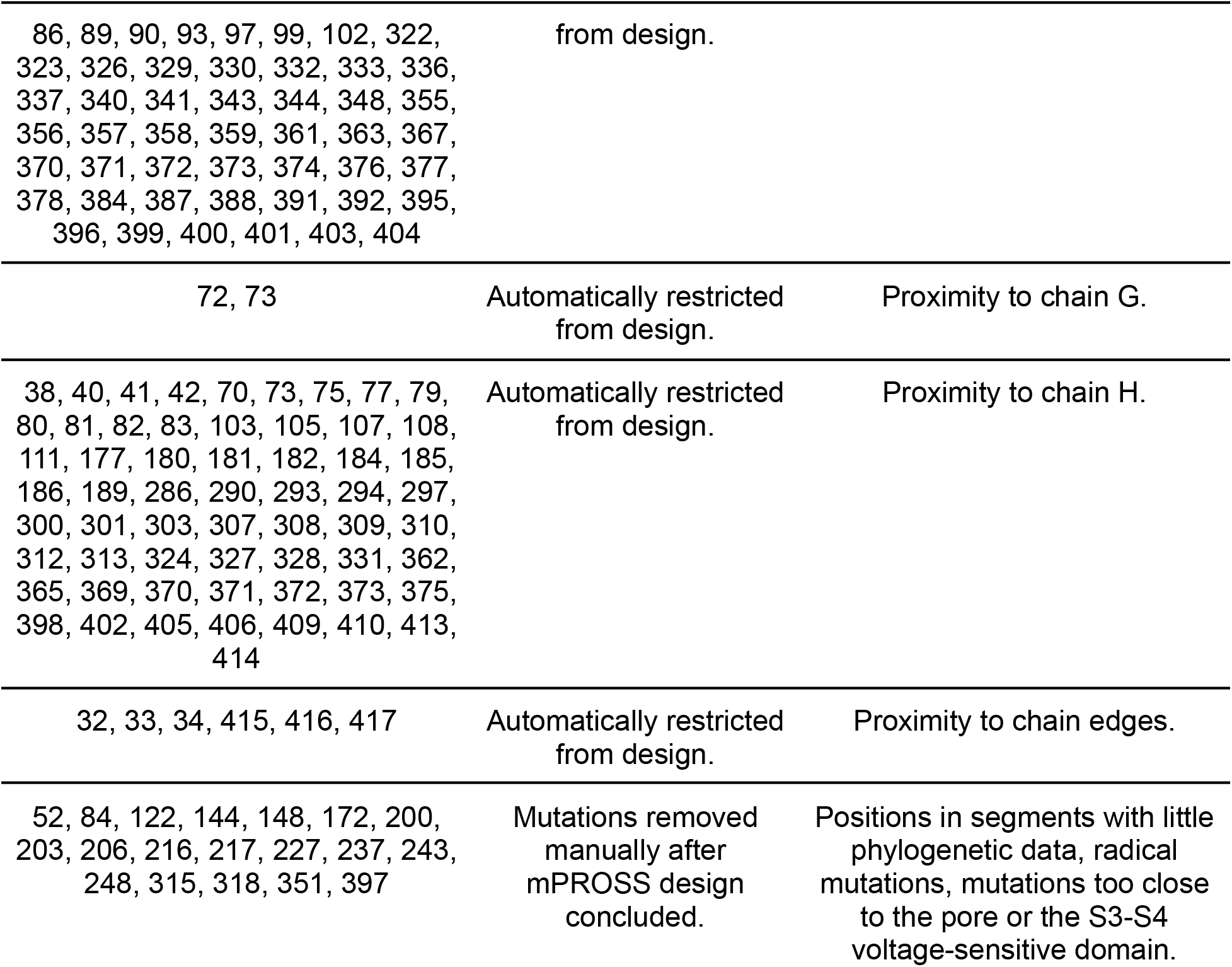
Positions that were restricted from design. * Positions are numbered the same as in the 2R9R PDB.

**Supplementary Table 4.**
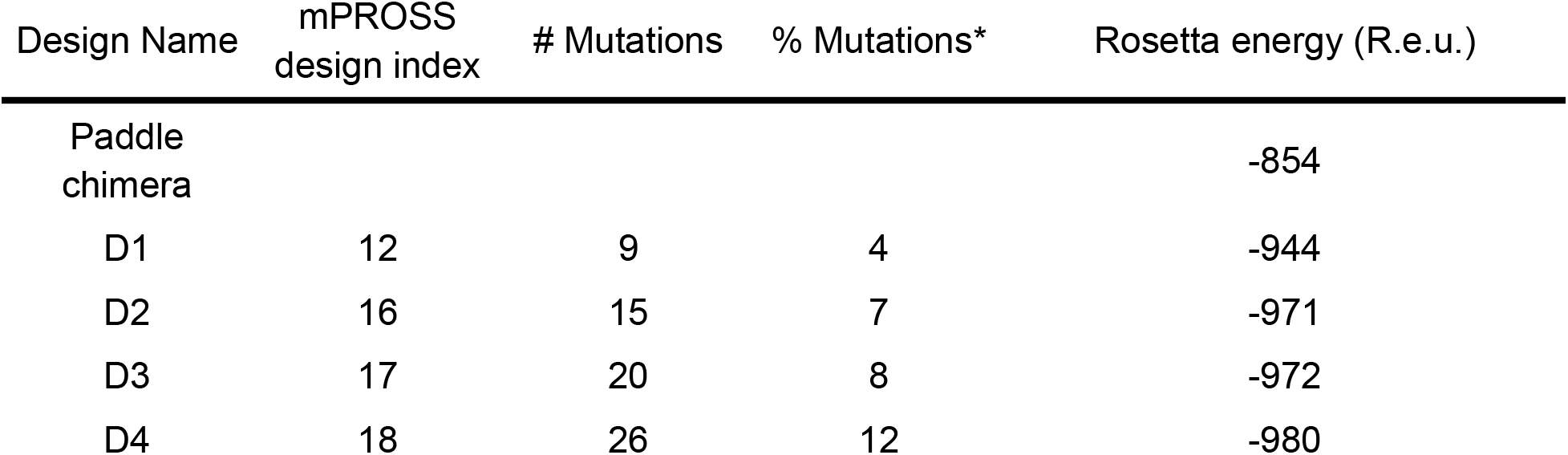
Design mutational load. * Percent of all designable positions (223 positions, which were resolved in the 2R9R PDB structure and were not restricted from design). R.e.u. — Rosetta energy units.

**Supplementary Table 5.** Mutated positions of interest. * Numbering according to Long et al.(27).

| 2R9R<br>position | Long et al.<br>numbering* | paddle chimera | D1 | D2 |
| --- | --- | --- | --- | --- |
| 56 | 75 | E | E | D |
| 114 | 133 | R | R | K |
| 132 | 151 | Y | Y | F |
| 242 | 261 | F | F | L |
| 252 | 271 | T | R | R |
| 261 | 280 | V | V | L |
| 321 | 340 | M | L | L |

**Supplementary Table 6:** Mean currents and V1/2 from oocytes expressing paddle chimera Kv1.2-Kv2.1 paddle chimera and various derivatives (n≥10). To account for variability between experiments, results from separate experiments were normalized relative to the paddle chimera. * Normalized to measurements of the paddle chimera from the same experiment.

| Construct | Normalized<br>current* | Normalized<br>$V_{1/2}$* |
| --- | --- | --- |
| Non-injected | 0.08±0.05 | 0.0±0.0 |
| Wild-type | 1.0±0.21 | 1.0±0.33 |
| D1 | 5.11±1.67 | -0.52±0.22 |
| D2 | 13.74±1.79 | -1.03±0.17 |
| D3 | 4.95±3.39 |  |
| D4 | 10.91±8.60 |  |
| D2 75E | 26.98±6.52 | -0.89±0.12 |
| D2 133R | 36.91±8.15 | -0.29±0.12 |
| D2 151Y | 33.55±4.2 | -1.07±0.12 |
| D2 261F | 12.42±3.68 | -1.1±0.26 |
| D2 280V | 27.1±2.58 | -1.55±0.49 |
| D2 271T | 25.16±4.11 | -0.92±0.24 |
| D2 340M | 45.67±7.47 | -0.68±0.24 |

**Supplementary File 1.**
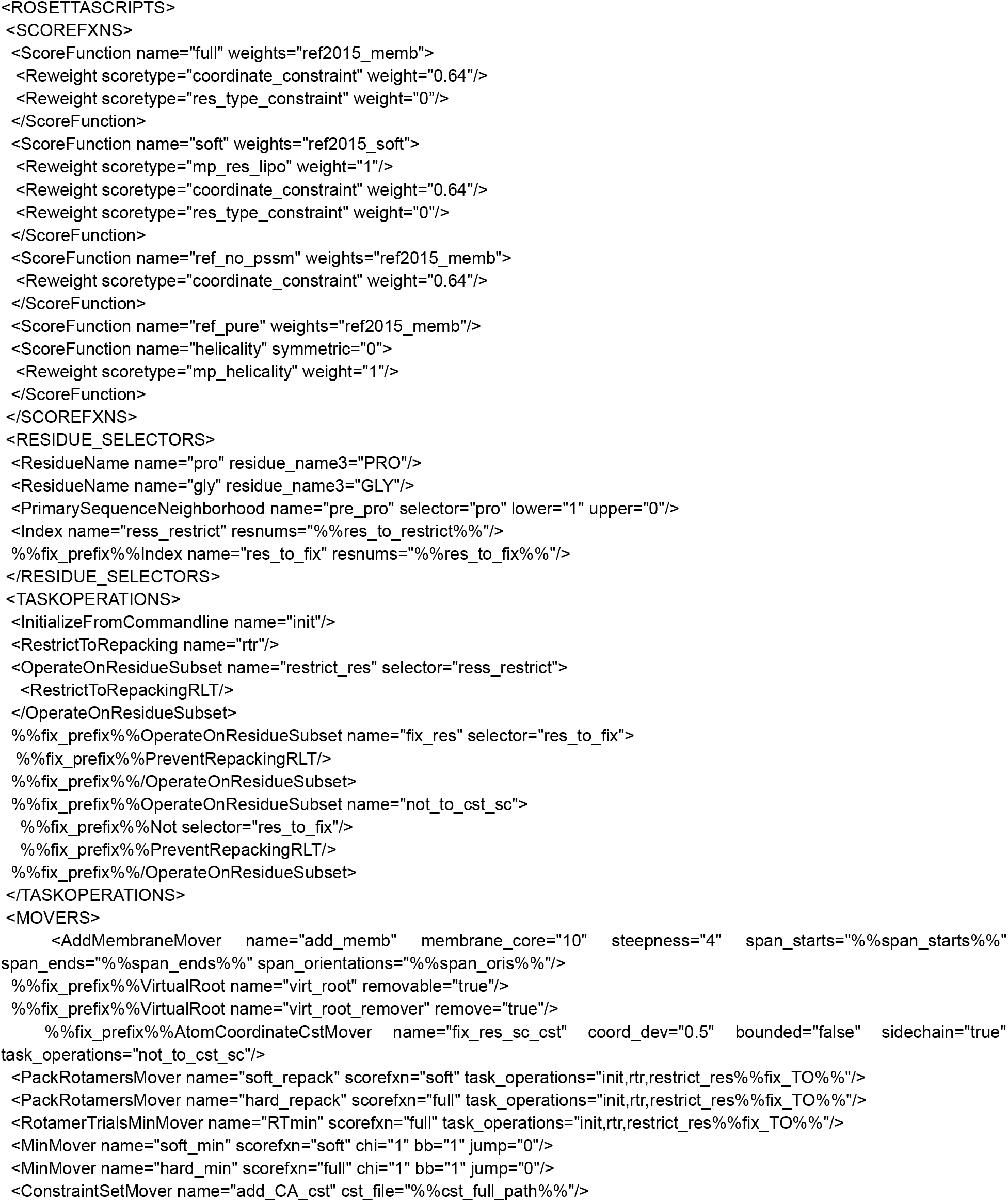
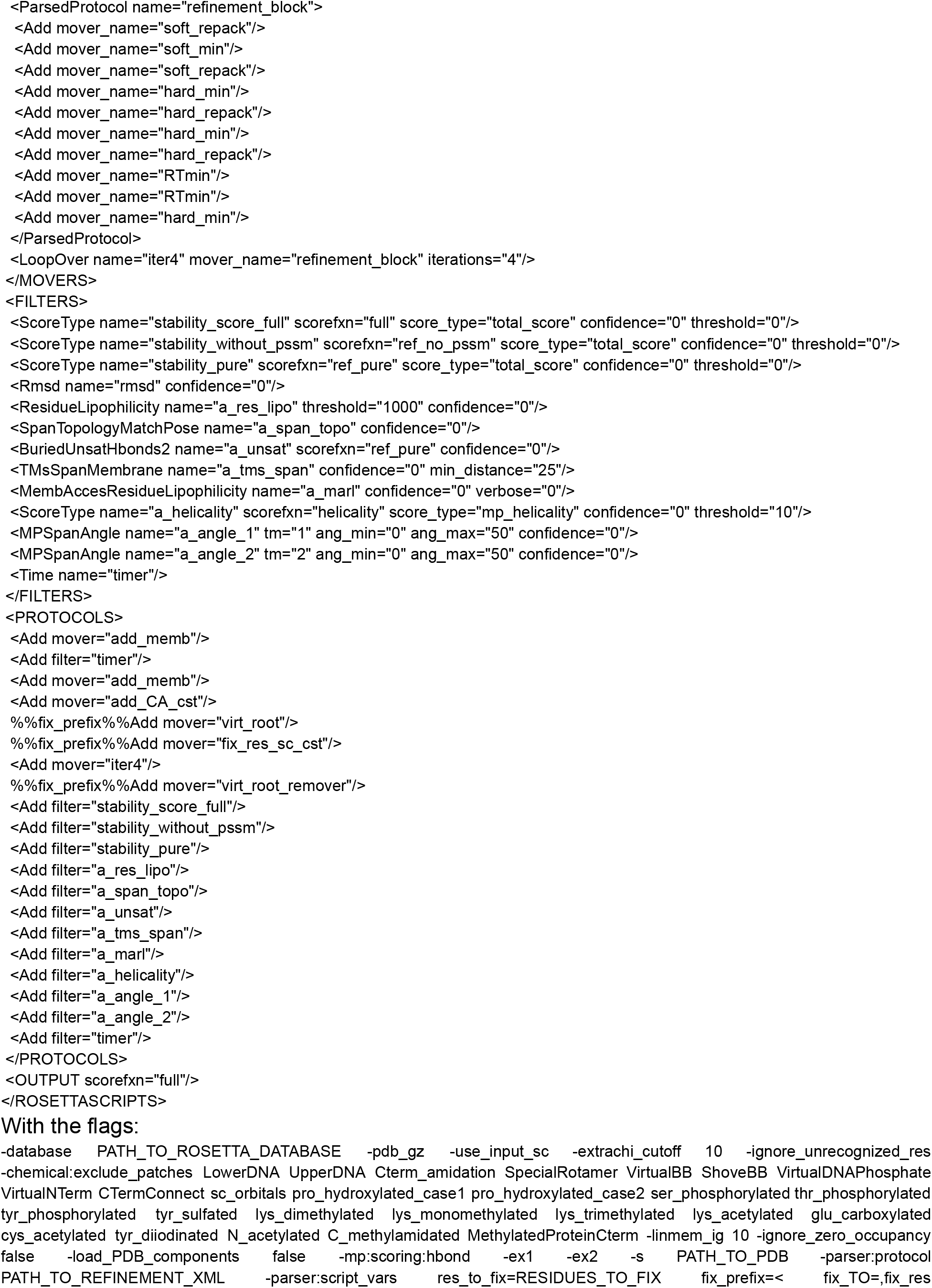
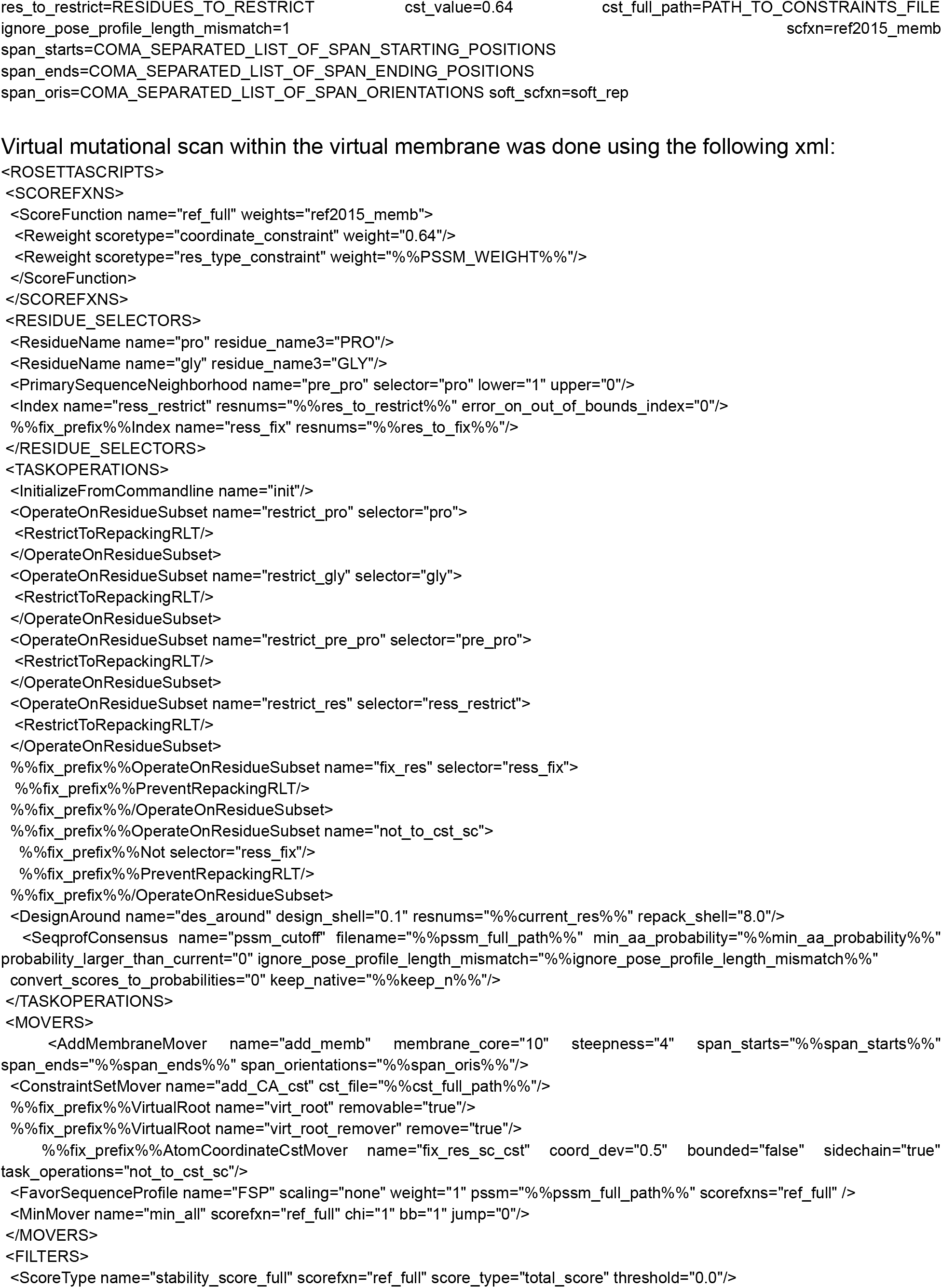
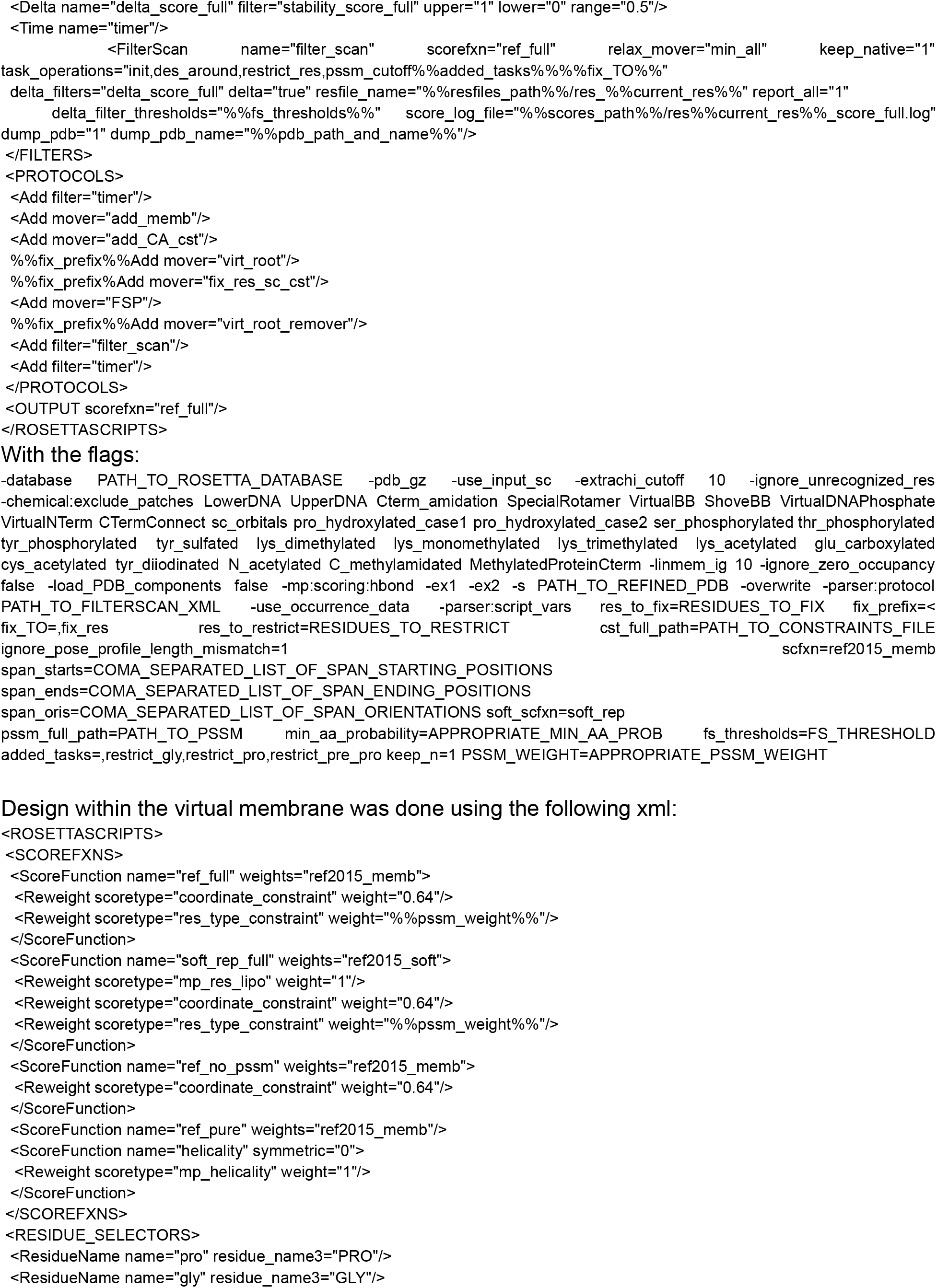
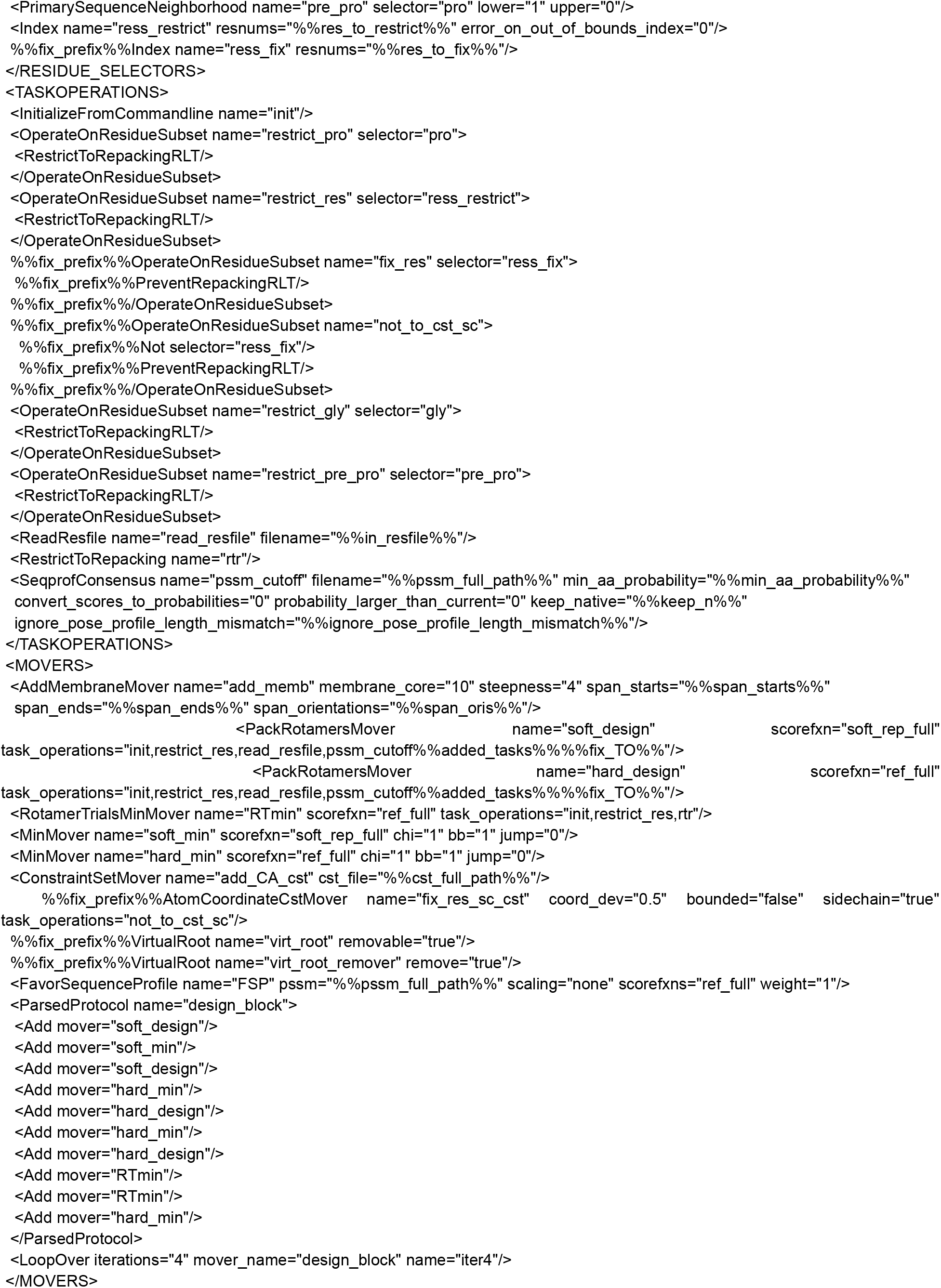
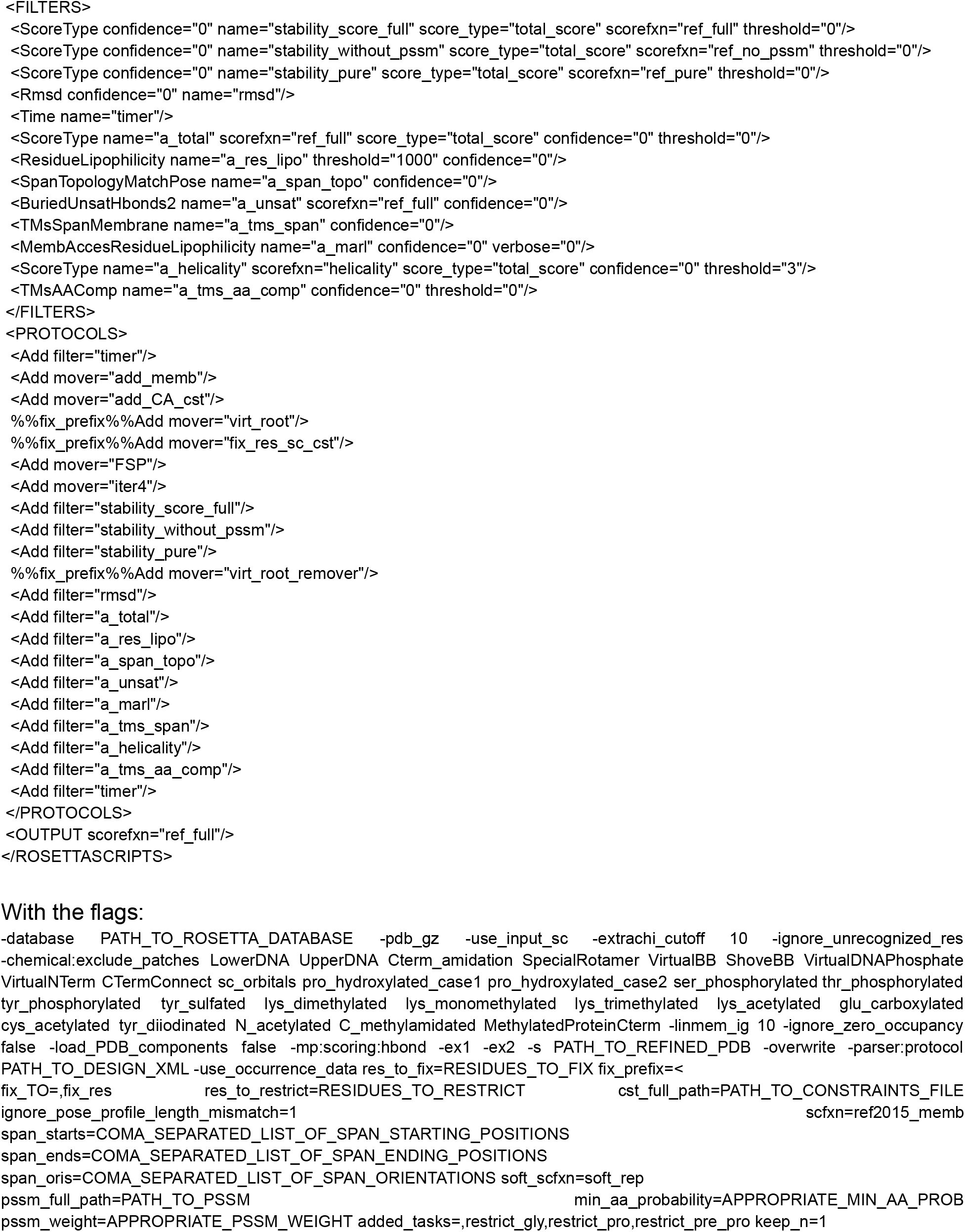
RosettaScripts scripts and flags for refinement, mutational scan, and design. All Rosetta calculations were done using the git commit b210d6d5a0c21208f4f874f62b2909f926379c0f. Refinement within the virtual membrane was done using the following xml:

**Supplementary File 2.**
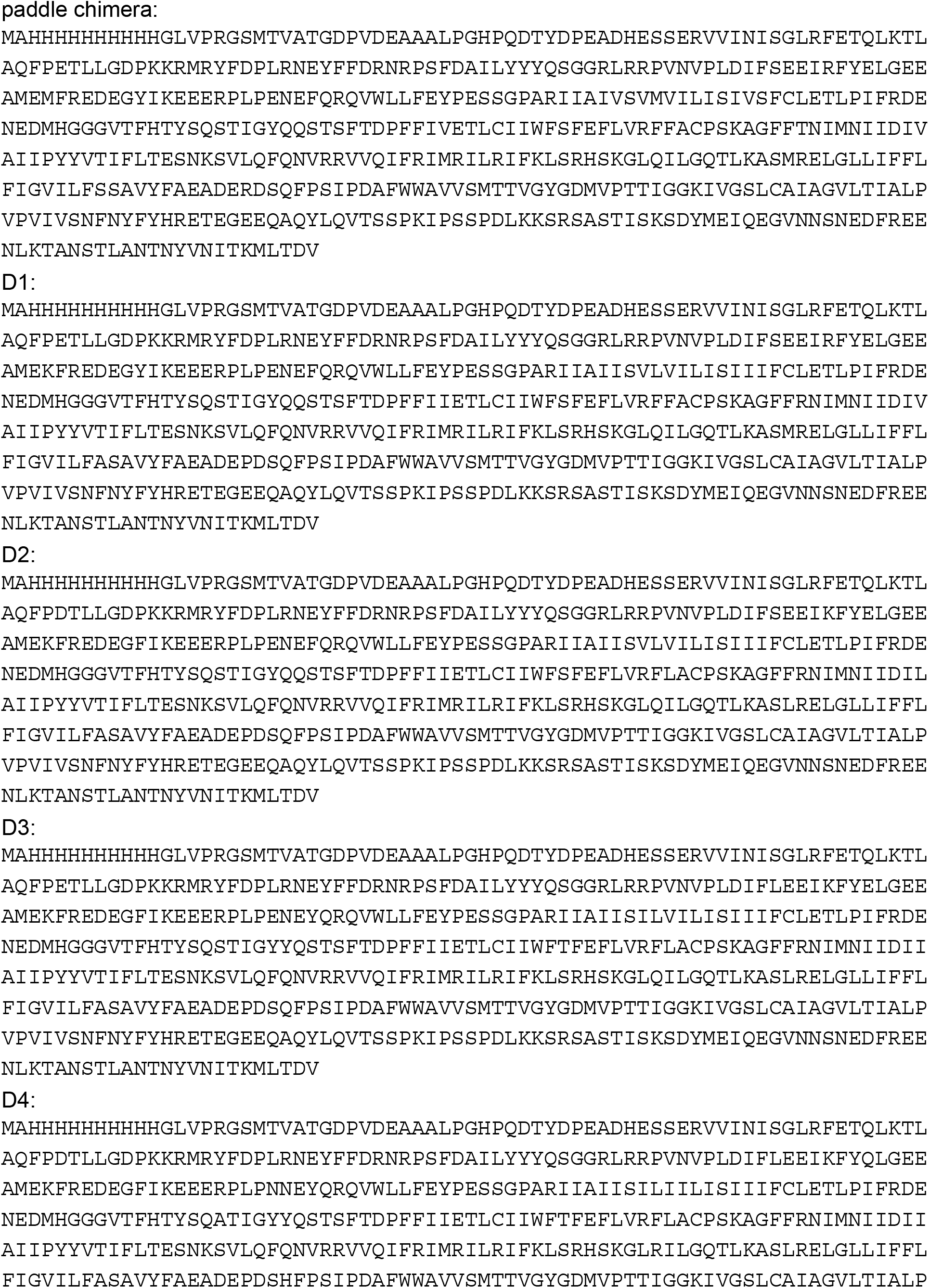
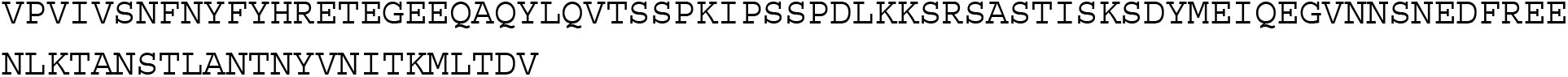
paddle chimera design sequences.

